# Lymphangiocrine signals are required for proper intestinal repair after cytotoxic injury

**DOI:** 10.1101/2022.03.08.479995

**Authors:** Brisa Palikuqi, Jeremie Rispal, Deedepya Vaka, Dario Boffelli, Ophir Klein

## Abstract

The intestinal epithelium undergoes continuous renewal and has an exceptional capacity to regenerate after injury. Maintenance and proliferation of intestinal stem cells (ISCs) is regulated by their surrounding niche, largely through Wnt signaling. However, it remains undetermined which niche cells produce signals during different injury states, and the role of endothelial cells (ECs) as a component of the ISC niche during homeostasis and after injury has been underappreciated. Here, we show that lymphatic endothelial cells (LECs) reside in proximity to crypt epithelial cells and secrete molecules that support epithelial renewal and repair. The LECs are an essential source of Wnt signaling in the small intestine, as loss of LEC-derived *Rspo3* leads to a lower number of stem and progenitor cells and hinders recovery after cytotoxic injury. Together, our findings identify LECs as an essential niche component for optimal intestinal recovery after cytotoxic injury.

## Introduction

Enteric blood and lymphatic networks are intimately associated with the small intestinal epithelium, including the ISCs that reside in epithelial invaginations called crypts (Bernier-Latmani et al., 2015; Bernier-Latmani and Petrova, 2017; Beumer and Clevers, 2020; Cifarelli and Eichmann, 2019). Over the last decade, several groups have shown that paracrine factors secreted by endothelial cells lining blood and lymphatic vessels, known as angiocrine factors, are important for stem cell maintenance and regeneration in many tissues (Augustin and Koh, 2017; Rafii et al., 2016). Blood endothelial cells (BECs) play a central role in the regeneration of several organs, such as liver, lung, bone marrow, and thymus (Butler et al., 2010; Ding et al., 2010; Ding et al., 2011; Ding et al., 2012; Hu et al., 2014; Wertheimer et al., 2018). Notably, LECs, through the secretion of the lymphangiocrine factor Reelin (RELN), have been linked to heart regeneration (Liu et al., 2020). In addition, the interaction of stem cells with LECs regulates stem cell cycling in the skin (Gur-Cohen et al., 2019). However, the role of intestinal ECs in ISC self-renewal and regeneration remains undetermined.

ISCs, which are marked by *Lgr5*, and their transit amplifying descendants self-renew and differentiate along the various intestinal epithelial lineages (Barker et al., 2007; Gehart and Clevers, 2019). Maintenance of ISCs is dependent on the niche, which is comprised of several cell types and is an essential source of Wnt ligands and modulators (McCarthy et al., 2020a; Palikuqi et al., 2021). Among these molecules, R-spondins (RSPOs), which bind to LGR receptors (LGR4 and LGR5 in the intestine), potentiate Wnt signaling (de Lau et al., 2011; Glinka et al., 2011) by stabilizing Frizzled receptors (Hao et al., 2012). RSPOs are essential for the expansion and renewal of ISCs both *in vivo* (Yan et al., 2017) and *in vitro* (Sato et al., 2009). RSPO3 is the predominant RSPO in the small intestine (Ogasawara et al., 2018), and inhibition of RSPO3 along with RSPO2 with a neutralizing antibody negatively affects epithelial regeneration after irradiation (Storm et al., 2015).

Several mesenchymal populations have been reported to be essential sources of Wnt ligands and modulators in the intestine (Degirmenci et al., 2018; Kabiri, 2014; McCarthy et al., 2020b; Stzepourginski et al., 2017; Valenta et al., 2016). For instance, mesenchymal cells marked by *Foxl1*^+^, known as telocytes, express high levels of *Wnt2* and *Wnt5a* (Aoki et al., 2016; Kondo et al., 2019; Shoshkes-Carmel et al., 2018). The inhibition of Wnt ligand secretion by deletion of *Porcupine* (*Porcn)* in *Foxl1*^+^ telocytes results in rapid stem and progenitor cell loss (Shoshkes-Carmel *et al.*, 2018). Another subtype of mesenchymal cells, known as trophocytes and marked by expression of *Grem1* and *Cd81* and low expression of *Pdgfra*, resides in proximity to ISCs and expresses the Wnt-family member *Wnt2b* as well as high levels of all R-spondins. Ablation of the trophocyte mesenchymal population leads to loss of *Lgr5*^+^ cells, while the population of transient amplifying cells remains stable (McCarthy *et al.*, 2020b). Moreover, deletion of *Rspo3* in *Pdgfra*^+^ cells does not have a major effect on epithelial cells during homeostasis (Greicius et al., 2018), suggesting that this essential signal is produced by several different niche cell types.

LECs express high levels of *Rspo3* (Kalucka et al., 2020; Ogasawara *et al.*, 2018), but it is unknown if LECs are an essential source of Wnt signaling in the small intestine during homeostasis or injury. Additionally, while several lines of evidence suggest that endothelial cells become activated and start proliferating in response to intestinal injury to drive epithelial recovery (Abel et al., 2005; Kinchen et al., 2018; Paris et al., 2001; Rehal et al., 2018), the contribution of endothelial cells and specifically LECs to intestinal maintenance and repair remains an open question. Here, we establish the spatial organization of lymphatic vessels in relation to crypt cells and utilize single cell RNA-sequencing (scRNAseq) to uncover the transcriptional composition of LECs, both during homeostasis and after cytotoxic injury. Our finding using mouse genetics that LEC-secreted *Rspo3* is essential for optimal intestinal repair after cytotoxic injury positions LECs as a key component of the intestinal niche and a critical source of Wnt signaling.

## Results

### Lymphatic endothelial cells reside in proximity to crypt cells

We first set out to delineate the location and molecular composition of LECs in the small intestinal niche. We utilized the CUBIC clearing method (Matsumoto et al., 2019), paired with whole mount imaging, to determine the spatial distribution of LECs in the mouse small intestine during homeostasis and to establish the relationship of LECs to BECs and mesenchymal cells in the niche. VECAD^+^ BECs and PDGFRA^+^ mesenchymal cells surrounded every crypt (**Figures 1A and S1B**), but lymphatic vessels displayed a unique pattern of organization. We found two distinct sets of lymphatic vessels: large lymphatic vessels that encircled the base of the crypts and gave rise to narrow lymphatic vessels in the villi, known as lacteals (**Figures 1A-B and S1A-C**). To determine the proximity of LECs to *Lgr5*^+^ stem cells in the crypts, we performed whole mount imaging on cleared small intestinal tissue from *Lgr5*^GFP-CreER^ mice. LECs are in proximity to *Lgr5*^*+*^ crypt cells at the base of the crypts and at the mid-crypt level (**Figures 1C and S1D**). More than 80 percent of crypts are in proximity to lymphatics at the base and/or mid-crypt levels, with 36 percent of crypts in proximity to LECs only at the base, 23 percent only at the mid-crypt region, and 26 percent at both (**Figure 1D**).

**Figure 1.**
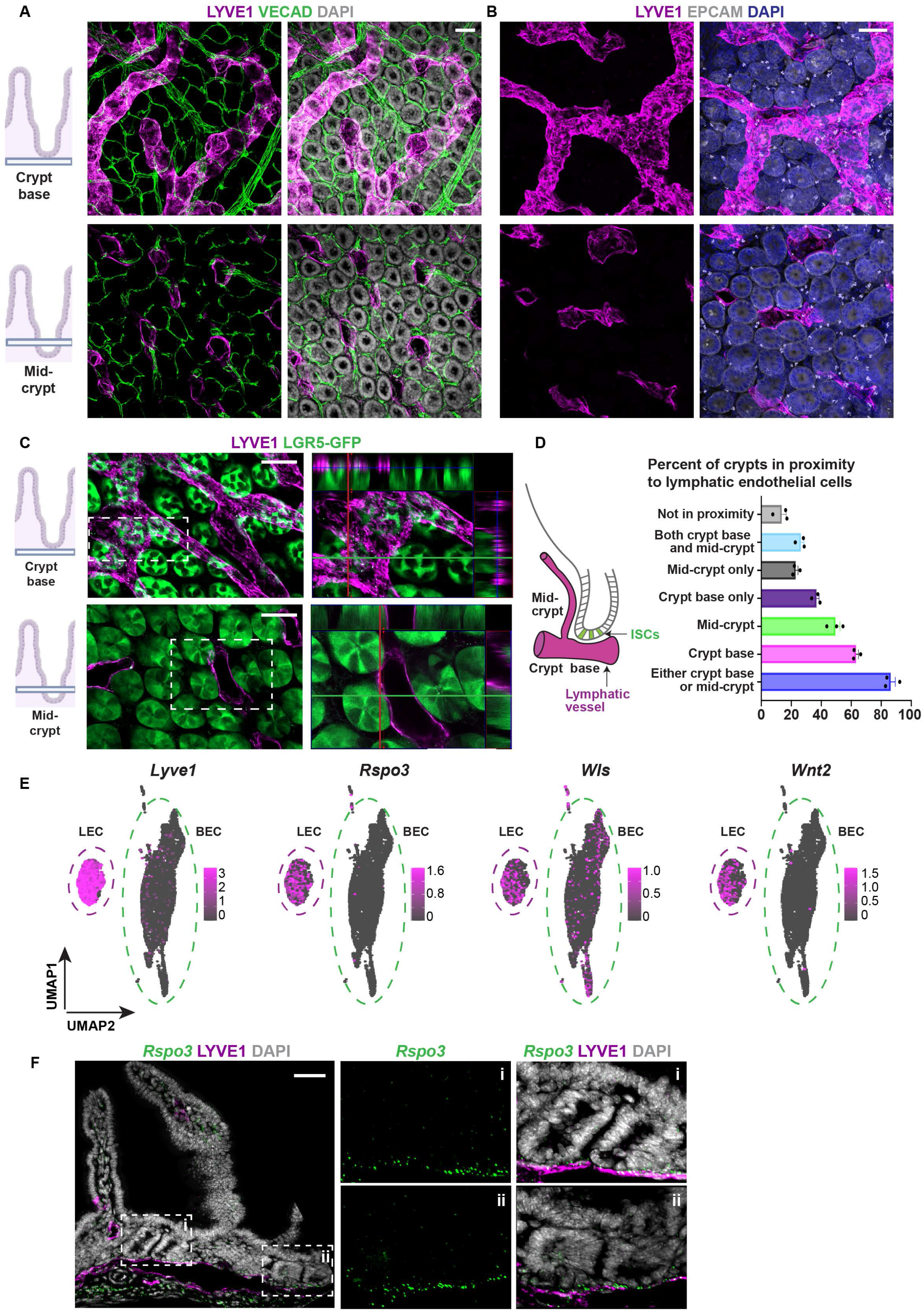
LECs reside in proximity to crypt cells and express high levels of *Rspo3*. **A-B)** Whole mount imaging of CUBIC cleared tissue from adult wild type mouse small intestine showing **(A)** organization of LECs and BECs at both the crypt base region and mid-crypt region and **(B)** large caliber lymphatic vessels residing in proximity to the base of the epithelial (*EPCAM*^*+*^) crypts cells and giving rise to smaller lymphatic vessels (lacteals). **C)** Whole mount imaging of CUBIC cleared tissue from adult *Lgr5*^GFP-Cre^ mice. LECs reside in proximity to *Lgr5*^*+*^ ISCs. Insets represent orthogonal projections of the whole mounts depicting adjacency and at times contact of LECs to *Lgr5*+ ISCs. **D)** Left: schematic representation of position of lymphatic vessels in relation to crypts. Right: quantification of percentage of crypts in vicinity to LECs at the base of the crypts, at mid-crypt level or not in proximity to any LECs (n=3 mice). **E)** scRNAseq analysis of ECs from wild type adult mouse small intestine. LECs (*Lyve1*^*+*^) in the small intestine express high levels of *Rspo3* and *Wls and Wnt2*. Data are from 4 mice over 2 independent experiments. **F)***In situ* hybridization (RNAscope) of adult mouse small intestine demonstrating expression of *Rspo3* mRNA in LECs that are in proximity to the crypt region. All scale bars = 50 μm. Data are mean +/− s.e.m.

### Lymphatic endothelial cells express high levels of *Rspo3*

Next, we set out to establish the molecular composition of LECs in the small intestine by performing scRNAseq. BECs and LECs were enriched by fluorescence-activated cell sorting (FACS) from the small intestine of wild type mice and identified as CDH5^+^, PODOPLANIN (PDPN)^Neg^ and CDH5^+^, PDPN^+^ cells, respectively (**Figures S2A and S2B**). In all, 6000 BECs and 2000 LECs were captured by scRNAseq. LECs clustered into two main groups, most likely representing LECs from the large vessels at the base of the crypt and lacteals (**Figures 1E and S2C**). LECs could also be distinguished from BECs by the high expression of *Aqp1, Cyp4b1, Il33 and Mmrn1* genes (**Figure S2E**). The transcription factor *Maf* is also selectively expressed in LECs (**Figure S2E**). Reelin (RELN), which is expressed in the heart by LECs and has been found to be essential for heart regeneration, is also highly expressed by LECs but not BECs in the small intestine (**Figure S2E**).

As Wnt signaling is essential for ISC maintenance and proliferation, we mined the scRNAseq data for the expression of Wnt-family members. As previously reported (Kalucka et al., 2020; Ogasawara et al., 2018), LECs in the small intestine expressed high levels of *Rspo3* (**Figures 1E and S2F**). Additionally, LECs expressed high levels of *Wntless* (*Wls*, essential for the secretion of Wnt ligands) and the canonical Wnt-family member *Wnt2* (**Figures 1E and S2F**). Interestingly, *Rspo3* and *Wnt2* were uniquely expressed by LECs, whereas *Wls* was also expressed by BECs (**Figure 1E and S2F**). *Rspo3* expression by LECs in proximity to epithelial crypt cells was confirmed by *in situ* hybridization (RNAscope) (**Figure 1F**); no other Wnt-family members were expressed by LECs during homeostasis.

### Loss of lymphatic *Rspo3* leads to impaired intestinal recovery after cytotoxic injury

As *Rspo3* is important for the proliferation of stem and progenitor cells in the crypt, we next utilized a genetic model to specifically delete *Rspo3* in LECs. We bred mice carrying the lymphatic driver *Prox1*^CreER^ with *Rspo3*^fl/fl^ mice to generate *Prox1*^CreERT2^;*Rspo3*^fl/fl^ mice (**Figure 2A**). Deletion of *Rspo3* in adult LECs did not significantly alter crypts in the small intestine during homeostasis. Control and knock-out mice had similar levels of EdU incorporation, crypt number and OLFM4 expression 2 weeks after tamoxifen induction (**Figure 2C-E**). Thus, during homeostasis, Wnt signaling from other, redundant niche components, such as PDGFRA^+^ cells, is sufficient for ISC maintenance.

**Figure 2.**
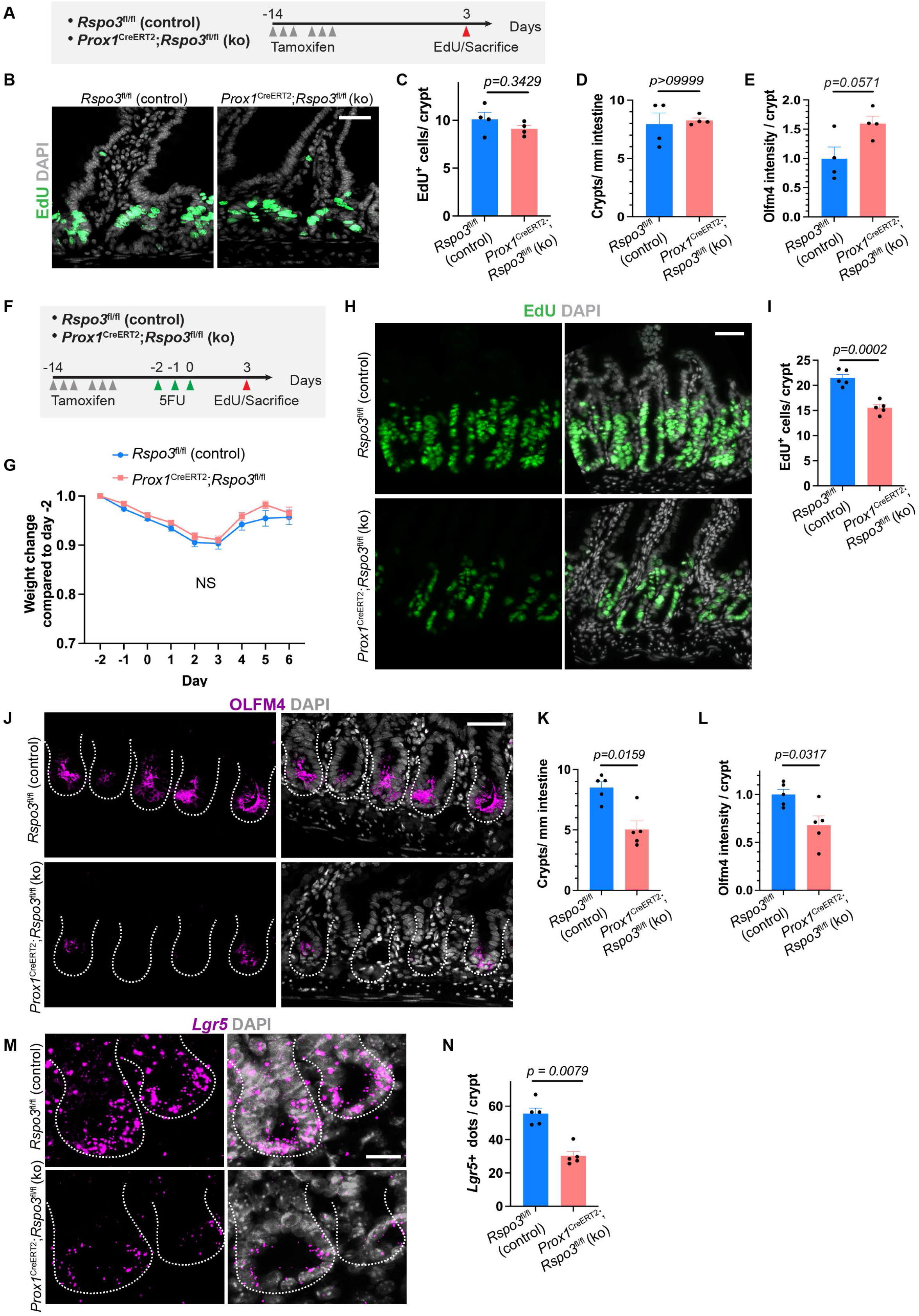
Lymphatic *Rspo3* is essential for intestinal recovery after 5FU injury. **A)** Tamoxifen induced *Prox1*^CreERT2^;*Rspo3*^fl/fl^ knock-out mice and control *Rspo3*^fl/fl^ were sacrificed 14 days after tamoxifen induction and after a 2-hour EdU pulse. **B)** Representative images of EdU staining from control and knock-out mice at homeostasis (n=4 mice per group) Scale bar = 50 μm. **C)** Quantification of EdU^+^ cells per crypt at homeostasis for control and knock-out mice. **D)** Quantification of total crypts per millimeter (mm) at homeostasis for control and knock-out mice. **E)** Quantification of OLFM4 staining intensity for control and knock-out mice at homeostasis. **F)** Control and *Rspo3* knock-out mice were treated for 3 days with 5FU at 75 mg/kg. Recovery was assessed at day 3 after injury. Mice were injected with EdU 2 hours before sacrifice. **G)** Weight change after 5FU injury in control and *Rspo3* knock-out mice compared to pre-injury (day −2). **H)** Representative images of EdU staining in the crypts of control and *Rspo3* knock-out mice at day 3 and **(I)** quantification of EdU^+^ cells per crypt at day 3 (n=5 mice per group). Scale bar = 50 μm. **J)** OLFM4 staining in control and knock mice at day 3. Scale bar = 50 μm. **K)** Quantification of OLFM4^+^ crypts/mm of small intestine at day 3. **L)** OLFM4 intensity staining per crypt at day 3. **M)** Representative images of *Lgr5+* in situ hybridization (RNAscope) for control and knock-out mice at day 3 and **(N)** quantification of *Lgr5*^*+*^ transcripts per crypt. Scale bar = 20 μm. Data are mean +/− s.e.m., Mann Whitney U-test, two-sided.

We then assessed whether lymphatic *Rspo3* is essential for intestinal recovery after injury, when crypt epithelial cells must undergo high proliferation. To test this hypothesis, we utilized 5-fluorouracil (5FU), a chemotherapeutic that selectively kills actively proliferating cells. Control and *Rspo3* knock-out mice underwent sublethal 5FU injury and were sacrificed at day 3 after the last 5FU injection (**Figure 2F-G**). After 5FU injury, whereas control crypt epithelium exhibited high levels of proliferation, knock-out mice had significantly decreased proliferation, as measured by EdU incorporation (**Figure 2H-I**). The total number of regenerating crypts, OLFM4^+^ cells, and *Lgr5*^+^ cells were also significantly reduced in knock-out mice (**Figure 2J-N**).

5FU treatment induces substantial damage to proliferating cells, potentially affecting both epithelial and niche cells. We therefore investigated whether a similar requirement for LEC-derived *Rspo3* exists when *Lgr5*^+^ cells alone are ablated. We generated *Prox1*^CreERT2^;*Rspo3*^fl/fl^;*Lgr5*^DTR^, in which tamoxifen injection leads to deletion of *Rspo3* in LECs, and Diphtheria Toxin (DT) (Tian et al., 2011) injection results in ablation of *Lgr5*^+^ cells. Tamoxifen induction was followed by *Lgr5*^+^ cell ablation for 3 consecutive days (**Figure S3A**). Although there was a slight decrease in levels of proliferation as measured by EdU incorporation, the effect was less pronounced compared with that observed after cytotoxic injury (**Figures S3B-F**). Thus, LEC-derived *Rspo3* is essential for adequate intestinal recovery when there is a requirement for a marked increase in proliferation of epithelial cells and when the niche becomes activated during injury.

### Loss of lymphatic *Rspo3* results in loss of transient amplifying and *Sca1*^*high*^ cells after cytotoxic injury

Next, we set out to uncover the molecular mechanism that hinders recovery in *Prox1*^CreERT2^;*Rspo3*^fl/fl^ knock-out mice after 5FU injury. At day 3 after the final 5FU injection, we isolated, from both control and knock-out mice, crypt epithelial cells as EPCAM^+^, CD44^+^; LECs as CD31^+^, PDPN^+^; BECs as CD31^+^, PDPN^neg^; and other stromal cells as EPCAM^neg^, CD31^neg^, and we then performed scRNAseq (**Figure 3A and S4A-B**). Analysis included 20,803 cells for control mice and 9,153 for knock-out mice, and all the merged samples grouped in 28 clusters (**Figure 3B and S4C**). Crypt epithelial cells were identified as *Epcam*^+^, *Cd44*^+^, and LECs were identified by the pan-endothelial markers *Cd31*, *Cdh5* or *Kdr* and expression of *Lyve1* or *Pdpn*. We also identified clusters of BECs as positive for *Cd31*, *Cdh5* or *Kdr* and negative for *Lyve1 and Pdpn*, mesenchymal cells as *Pdgfra*^+^, and myofibroblasts as *Myh11*^+^ (**Figure 3C-D and S4D**). Within the *Pdgfra*^+^ mesenchymal populations, we identified telocytes as *Foxl1*^+^, trophocytes as *Cd81*^+^, *Pdgfra*^low^, and pericryptal stromal cells as *Cd34*^*+*^ (**Figure S4E-F**).

**Figure 3.**
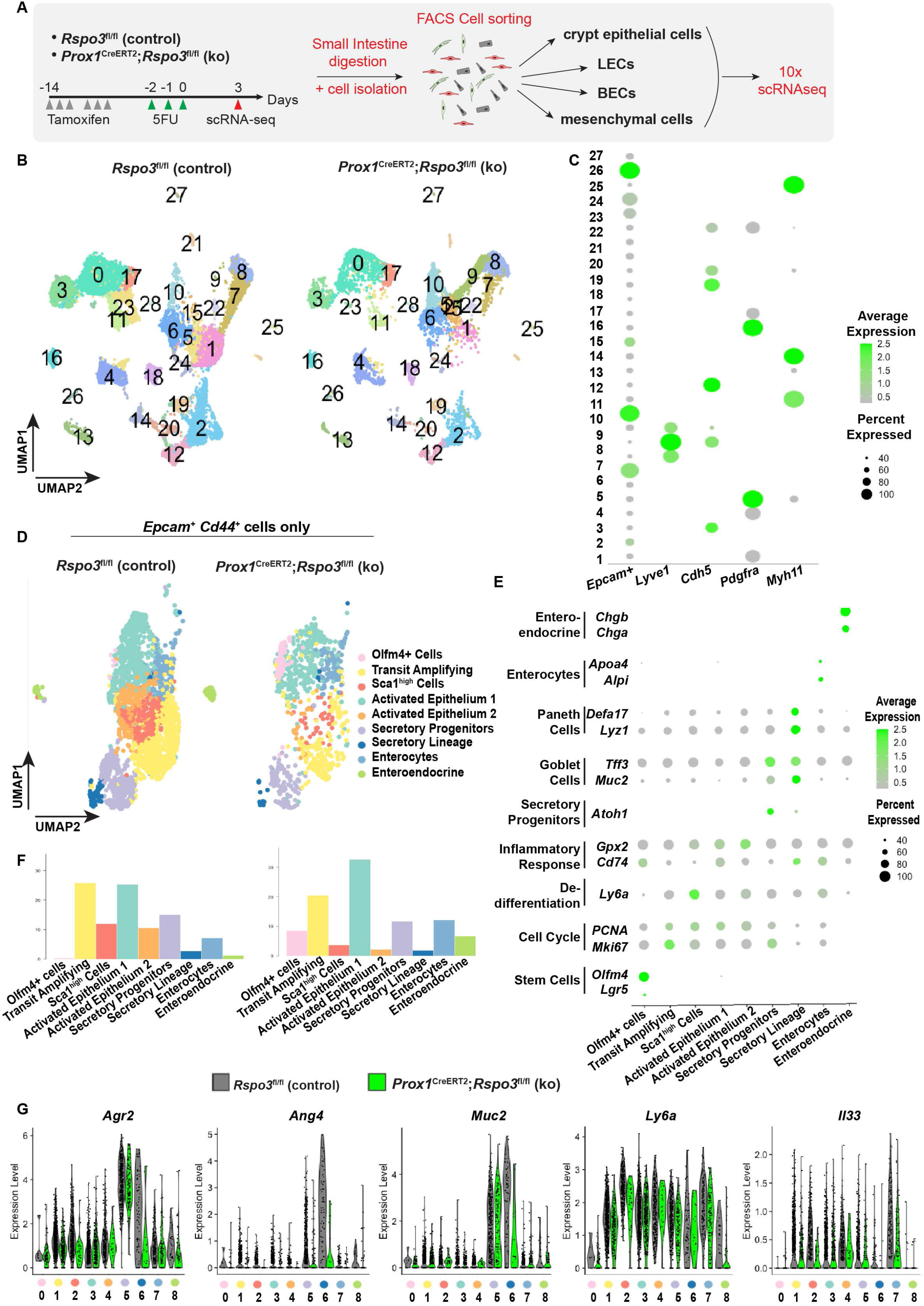
scRNAseq reveals that loss of lymphatic *Rspo3* results in reduced number of TA and *Sca1*^*high*^ cells after cytotoxic injury. **A)** Schema of scRNAseq experiment. Tamoxifen treated control and knock-out mice were administered 5FU on 3 consecutive days. Crypt epithelial cells, LECs, BECs and other stromal cells were enriched by FACS, and the samples submitted for 10X scRNAseq analysis. Data are from 2 independent biological experiments. Each biological experiment included 2 control and 2 knock-out mice. **B)** The sequenced cells group in 28 clusters as depicted in UMAP (Uniform Manifold Approximation and Projection) plots. **C)** Dotplot of main markers and corresponding clusters for LECs (*Lyve1*^*+*^), epithelial cells (*Epcam*^+^), BECs (*Cdh5*^+^), mesenchymal cells (*Pdgfra*^+^), and myofibroblasts (*Myh11*^+^). **D)** Crypt epithelial cells were identified as *Epcam*^*+*^ *Cd44*^*+*^ cells and re-clustered. Crypt epithelial cells group in 9 distinct clusters as shown in UMAP plots. **E)**Dotplot of main markers identifying the 9 clusters represented in *Epcam*^*+*^ *Cd44*^*+*^ cells. **F)** Bar graph of the proportional distribution of cells within crypt epithelial clusters. **G)**Violin plot of differentially expressed genes (DEGs) in crypt epithelial cells from control (grey) versus knock-out (green) mice.

To compare the effect of 5FU treatment on crypt epithelial cells in control and knock-out mice, we isolated *Epcam*^+^ *Cd44*^+^ cells from the merged cell population and re-performed cluster analysis (**Figure 3E**). We identified 9 clusters in *Epcam*^+^ *Cd44*^+^ epithelial cells: stem cells (*Olfm4*^+^, *Lgr5*^+^), cycling transit amplifying cells (TA) (*Mki67*^+^), secretory progenitors (*Atoh1*^+^), secretory cells (including Paneth cells as *Lyz1*^+^ and goblet cells as *Muc2*^+^), enteroendocrine cells (*ChgA*^+^, *ChgB*^+^) and mature enterocytes (*Alpi*^+^) (**Figures 3D and 3E**). We also identified 3 other clusters of crypt epithelial cells. One was a *Sca1*^high^ (*Ly6a*) cluster; *Sca1* has been identified as a marker of dedifferentiation and transition to a fetal-like state in the small intestine after injury (Nusse et al., 2018). We also identified 2 clusters with elevated expression of markers such as *Cd74* and *Gpx2* (activated epithelium 1 and 2), which have been associated with activated epithelium in bacterial and parasitic infections (Haber et al., 2017) (**Figures 3D-E**).

We determined differential cell composition and gene expression between control and knock-out samples for each cluster. Control crypt epithelial cells had a higher proportion of TA and *Sca1^high^* cells (**Figure 3F**), and mutant crypt epithelial cells had lower expression of *Agr2*, *Ang4*, *Mu*c2, *Ly6a* (*Sca1*) and *Il33*, amongst others (**Figure 3G**). *Agr2* is important for mucus production and processing, and its loss has been associated with mucus barrier dysfunction and colitis (Zheng et al., 2006). *Ang4* is an antimicrobial peptide involved in innate immunity and epithelial host defense in the small intestine (Forman et al., 2012; Hooper et al., 2003). Lower numbers of TA and *Sca1*^*high*^ cells in mutant mice than in control were accompanied by a higher proportion of *Olfm4*^+^, *Lgr5*^+^ cells compared to other crypt epithelial cells (**Figure 3G**). Thus, loss of *Rspo3* from LECs leads to decreased numbers of TA and *Sca1*^high^ cells, and to decreased mucus antimicrobial peptides genes, resulting in an overall impairment of the recovery process of crypt epithelial cells after 5FU injury in the absence of LEC-derived *Rspo3*.

### The immunomodulatory and inflammatory response to 5FU injury is repressed in *Rspo3* knock-out LECs

Next, we investigated changes in LECs after 5FU injury in control versus knock-out mice. We performed whole mount imaging at day 3 after the last 5FU injection. LEC density was found to be similar in control and knock-out mice (**Figure 4A-B**). In order to identify transcriptional changes in LECs after 5FU injury, we further isolated endothelial cells as *Cdh5*^+^, *Kdr*^+^ or *Cd31*^+^ from our scRNAseq data and performed cluster analysis. All endothelial cells grouped into 11 distinct clusters (**Figure 4C**). Amongst those, LECs were identified by the expression of *Lyve1* and included 2 clusters: cluster 0 and 3 (**Figure 4C-D**). Interestingly, cluster 3 consisted almost exclusively of knock-out LECs after 5FU damage (**Figure 4C**). In order to better understand these transcriptional changes in LECs, we performed differential expression analysis. Pathway analysis of genes differentially upregulated in control LECs revealed an enrichment of pathways related to the immune and inflammatory response of LECs. Inflammatory genes such as *Cebpb* (Ruffell et al., 2009) and *Mif* (Abu El-Asrar et al., 2019) and anti-microbial genes such as *Defa24*, *Defa30*, *Lyz1, and Agr2* were expressed at significantly higher levels in control LECs compared to knock-out LECs (**Figure 4G**). Conversely, genes related to endothelial-mesenchymal transition such as *Prss23, (Bayoumi et al., 2017), Tgfb1, Tgfb2 (Maleszewska et al., 2013; Yoshimatsu et al., 2020)* and *Vim*, and those shown to have anti-lymphangiogenic effects such as *Cavin1 (Yang et al., 2020)*, were expressed at significantly higher levels in *Rspo3* knock-out LECs (**Figure 4H**). Cluster 3, which consists largely of knock-out LECs, was especially enriched in these genes (**Figure 4C,4H**). In sum, loss of *Rspo3* from LECs leads to reduced crypt proliferation after cytotoxic injury, resulting in a lower number of TA and *Sca1*^high^ crypt epithelial cells and a dampened immunomodulatory response in LECs.

**Figure 4.**
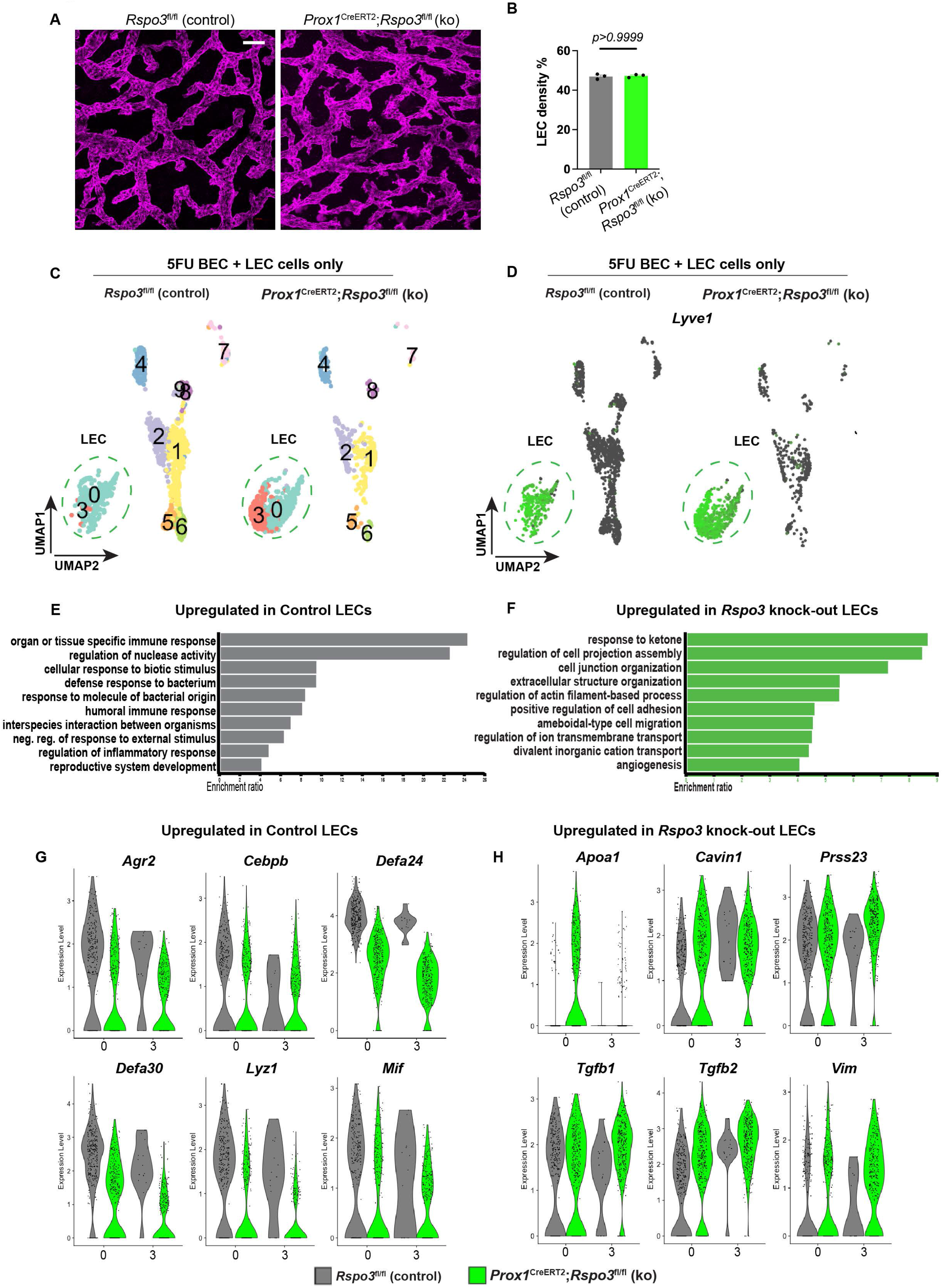
Deletion of *Rspo3* in LECs hinders the expression of immune supporting genes after 5FU injury. **A-B)** Whole mount images of control and knock-out mice at day 3 after 5FU treatment and quantification of LEC density at day 3 after 5FU treatment. **C)** ECs were identified as *Cdh5*^*+*^, *Cd31*^*+*^ or *Kdr*^*+*^ cells and re-clustered. ECs group in 10 distinct clusters. **D)** Clusters 0 and 3 were identified as lymphatic based on *Lyve1* expression. **E-F)**GO terms of biological pathways of enriched in genes upregulated in **(E)** control LECs **(F)** and knock-out LECs were identified by utilizing WEB-GESTALT. **H)** Violin plots of DEGs enriched in **(G)** control or **(H)** knock-out LECs; grey violin plots represent LECs from control mice and green violin plots represent LECs from knock-out mice. Data are mean +/− s.e.m., Mann Whitney U-test, two-sided.

## Discussion

The cellular components that constitute the ISC niche are currently the subject of intensive investigation, but the involvement of endothelial cells has been understudied. We report that LECs are physically part of the niche, with large vessels placed just under crypts and in proximity to ISCs, and with smaller lacteal vessels that ascend along the crypt to the top of villi. In all, over 80% of crypts are in proximity to lymphatic vessels. Additionally, analysis of our scRNAseq data revealed that LECs also secrete several components of the Wnt pathway, such as *Rspo3*, *Wls* and *Wnt2*. Deleting *Rspo3* specifically in LECs utilizing the *Prox1* driver demonstrated that LEC-secreted *Rspo3* is essential for the proper regeneration of the intestinal epithelium after injury caused by 5FU treatment.

Angiocrine factors support stem cells and regenerative processes of several tissues. This has led to an appreciation that ECs are more than plumbing that carries blood and oxygen, as they provide essential molecular signaling for maintaining tissue function. Our findings demonstrate that lymphangiocrine-derived factors in the intestine are critical for intestinal regeneration after cytotoxic injury. This suggests that LECs could be an essential source of secreted factors in other tissues as well.

*Rspo3* produced by LECs is essential to support regeneration after 5FU-induced injury, but it is dispensable during homeostasis. This suggests that *Rspo3* signaling is provided by several different sources in the ISC niche during homeostasis. Consistent with this notion, other studies have shown that deletion of *Rspo3* in fibroblasts (*Pdgfra*^+^ (Goto et al., 2022; Greicius *et al.*, 2018)) or in myofibroblasts (Harnack et al., 2019) alone, or generalized inhibition of RSPO3 by a neutralizing antibody (Storm et al., 2015), has no obvious effect on epithelial homeostasis. We report that after damage that requires high levels of proliferation, such as 5FU, *Rspo3* secreted by LECs is essential for the timely repair of epithelial cells. In addition, LECs are relatively unaffected by deleterious conditions, such as 5FU (this paper) or irradiation (Goto et al., 2022). It therefore appears that intestinal maintenance and regeneration rely on different niche sources of *Rspo3*, presumably depending on the level of damage and required proliferation to achieve complete repair.

The finding of a supportive role for LECs to ISCs and regenerative cells is also relevant to diseases in the gut. In colorectal cancer (CRC), lymphangiogenesis is one cause of the development of metastasis (Li et al., 2011; Tacconi et al., 2015). Our finding that after 5FU treatment LEC-secreted *Rspo3* is essential to support ISC maintenance suggests that LECs could also be involved in the promotion of cancer. Furthermore, maintenance of the lymphatic vessel density of LECs after 5FU treatment raises questions about the potential involvement of LECs in cancer survival and escape from chemotherapy.

## Acknowledgments

This work was funded by NIH 35-DE026602 and by U01DK103147 from the Intestinal Stem Cell Consortium, a collaborative research project funded by the National Institute of Diabetes and Digestive and Kidney Diseases and the National Institute of Allergy and Infectious Diseases, to O.D.K. B.P. was supported by UCSF CIRM Scholars Training Program EDUC4-12812. We thank F. de Sauvage for critical discussions, helpful comments, and reagents. We thank B. Hoehn, A. Rathnayake, E. Sandoval, K. Pan, P. Marangoni and A. Verma for technical assistance and animal maintenance. We thank V. Nguyen at the UCSF Parnassus Flow Cytometry Core for assistance with FACS experiments and A. Poon and C. Chu for assistance with 10x Chromium and library preparation.

## RESOURCE AVAILABILITY

### Lead contact

Further information and requests for resources and reagents should be directed to and will be fulfilled by the lead contact, Ophir Klein.

### Materials availability

All the materials will be available upon request to the lead contact under material transfer agreement with UCSF.

### Data availability

All scRNAseq has been deposited on GEO and will be available before publication.

## EXPERIMENTAL MODEL AND SUBJECT DETAILS

### Mouse strains

All mouse experiments were conducted under the IACUC protocol AN176864 at University of California San Francisco. All mice (male and female) were used at 8-12 weeks of age at the beginning of each experiment. *Rspo3*^*fl/fl*^ mice were a gift of Christof Niehrs (Scholz et al., 2016), *Prox1*^*CreERT2*^ mice were purchased from Jackson labs (022075), *Lgr5*^*DTR/GFP*^ were originally derived at Genentech and are a kind gift of Fred de Sauvage (Tian *et al.*, 2011), *Lgr5*^GFP-IRES-CreERT2^ were purchased from Jackson labs (008875; (Barker *et al.*, 2007)). For knock-out induction, mice were gavaged with tamoxifen in corn oil (200 mg/kg) 6 times, 2 x 3 days interspersed by 3 days of rest. For EdU proliferation experiments, mice were injected with EdU (5mg/mL) 2 hours before sacrifice. For injury experiments, mice were injected for 3 consecutive days with either 5FU at 75 mg/kg in 0.9% saline or Diphtheria toxin (DT) (50 μg/kg). For vessel visualization, mice were injected with an antibody against VECAD (25 μg/mouse in 100 μl of PBS) conjugated to Alexa-647 8 min prior to sacrifice.

## METHOD DETAILS

### Tissue collection and preparation for microscopy

For all microscopy experiments, the jejunum was collected and fixed in 4% PFA for 24h. For sections, tissues were incubated in 30% sucrose for 24h at 4°C and then embedded in OCT before being sectioned to a thickness of 8 μm. For whole mount, fixed tissues were cleared for 3 to 6 days in CUBIC-L solution (10% N-butyldiethanolamine, 10% Triton X-100 (Matsumoto et al., 2019).

### Immunofluorescence, RNAscope, EdU labeling on sections and Whole mount staining

Frozen sections were baked at 60°C for 30min, unmasked for 30min, blocked (5% NDS, 1X Animal Free blocking, 0.3% Triton X-100) for 1h and incubated overnight with primary antibodies at 4°C (table). Then, sections were incubated with secondary antibodies for 1h at RT (table).

RNAscope was done according to the manufacturer’s protocol. Briefly, the sections were post-fixed in 4% PFA, incubated with hydrogen peroxide to inhibit endogenous peroxidase before performing target retrieval. Subsequently, the tissues were incubated with specific RNA probes (table), followed by revelation and amplification steps.

For EdU labeling, after the baking, slides were permeabilized for 5min in 1% Triton X-100 and EdU staining performed according to manufacturer protocol (table).

For whole mount staining, 1 cm tissue pieces were blocked for 1 day, incubated with primary and then secondary antibodies for 2 days and finally post-fixed for 1 day in 4% PFA (Bernier-Latmani, 2016). Each step was done at 4°C.

For all staining and labeling, tissues were counterstained with DAPI and mounted in Prolong Gold. The images were taken using a Zeiss Apotome 910 system.

### Quantification of staining

All image quantifications were done using ImageJ software. The proliferative cell number was quantified by counting the number of EdU^+^ cells per crypt and calculating the mean for each mouse. The crypt number was calculated by counting OLFM4^+^ crypts in 20 to 30mm of tissue length. The OLFM4 intensity was analyzed by calculating the mean of the staining in a defined area corresponding to the crypt. *Lgr5* RNAscope was quantified by calculating the mean of positive dots in each crypt. For the quantification of crypt number in close proximity to lymphatic vessels, crypts were assigned to different categories: 1) crypt base: at least one cell of the crypt base is above a lymphatic vessel, 2) mid-crypt: at least one side of the mid-crypt is beside of a lymphatic vessel, 3) mid-crypt only, 4) crypt base only, 5) both crypt base and mid-crypt, 6) either crypt base or mid-crypt and 7) not in proximity: neither crypt-base nor mid-crypt. For the quantification of lymphatic vessel density, after Z-projection of all optical sections, the area fraction of the LYVE-1 signal (LYVE-1 positive area on total area) was quantified for each mouse.

### Single cell RNA sequencing experiment

Small intestines from adult male and female mice were collected and cut into small pieces (0.5 cm) and put in 30 mL of dissociation solution (RPMI + 1% Glutamax, 1% non-essential amino acids, 2.5% HEPES, 1% Pen-strep, 5mM EDTA, 3% FBS and 1 μM DTT) in a rotator at 37°C for 30 min. Tissues were washed twice in HBSS, spun down and further minced using scissors. The minced tissue was then digested in a rotator for 30 min at 37°C in 4 ml of digestion solution (Collagenase A (2.5 mg/ml), Dispase II (1 U/ml) and DNAse (30 μg/ml) in HBSS with 1% Glutamax, 1% non-essential amino acids and 2.5% HEPES). The cells were strained (100 μM strainer) and the left-over tissue digested for another 30 min in more 4 ml of digestion solution. Both fractions were stained with antibodies and Aria II with a 100-micron nozzle utilized to perform the sort. For homeostasis experiments: lymphatic endothelial cells were sorted as CD31^+^, PDPN^+^, EPCAM^neg^, CD45^neg^ and blood endothelial cells as CD31^+^, PDPN^neg^, EPCAM^neg^ and CD45^neg^. For 5FU experiments: at day 3 after the last 5FU injection, small intestines were collected from control and knock-out male and female mice (8-12 weeks). Crypt epithelial cells were sorted as EPCAM^+^, CD44^+^, CD45^neg^, CD31^neg^, lymphatic endothelial cells were sorted as CD31^+^, PDPN^+^, EPCAM^neg^, CD45^neg^, blood endothelial cells as CD31^+^, PDPN^neg^, EPCAM^neg^, CD45^neg^, epithelial cells as EPCAM^+^, CD45^neg^, CD31^neg^ and stroma cells EPCAM^neg^, CD45^neg^, CD31^neg^. After sorting, the different populations were mixed and loaded into 10x Chromium. Two mice were used for each condition and the whole experiment was repeated twice on two different days for both homeostasis and 5FU experiments.

### scRNAseq analysis

The single-cell suspension was loaded onto the 10X Chromium Single Cell instrument (10X Genomics). Single-cell barcoding, and cDNA libraries were constructed using the 10X Chromium single-cell 3’ Library Kit according to the manufacturer’s protocol. Libraries were sequenced on an Illumina NovaSeq 6000 and the sequence output processed with Cell Ranger 5.0.1 to produce filtered feature-barcode count matrices (http://10xgenomics.com).

Single-cell analyses were performed using the Seurat package in R (v.4.1.0) (Butler et al., 2018). Count matrices filtered to retain only genes expressed in three or more cells in a sample, were used for further analysis. Cells with high mitochondrial gene percentage or in the 1^st^ or 99^th^ percentile of transcript counts were also removed. Initially, all sequenced cell types were analyzed together. Data were processed following standard practices, including log-normalization of UMI counts, and regression of mitochondrial gene expression. Principal component analysis was subsequently performed on the most variable genes. Cluster identification (at a resolution of 0.5) and UMAP visualization was performed on the top 20 principal components. Differential gene expression for gene-marker discovery across the clusters was performed using the Wilcoxon rank-sum test.

Subsequently, for 5FU experiments the following analyses were performed. 1) For crypt epithelial cells: *Epcam*^*+*^, *Cd44*^*+*^ cells were identified as having expression of *Epcam* >1, *Cd44* >1 and expression of *Cdh5*, *Cd31*, *Kdr* and *Pdfgra* <1. *Myh11*^*+*^ cells were also excluded. 2) ECs were isolated as *Cdh5*, *Cd31*, *Kdr1* >1 and *Epcam*, *Cdh1*, *Pdgfra1* <1. EC clusters that expressed *Lyve1* were identified as LECs. 3) Mesenchymal cells were isolated as *Pdgfra*>1, *Cdh5*, *Kdr*, *Cd31*, *Epcam*, *Cdh1* <1. For each cell type, filtering, clustering, and differential gene expression testing was done as described above. For pathway analysis, WEB-GESTALT (WEB-based Gene SeT AnaLysis Toolkit) (Liao et al., 2019) was utilized, where genes that had at least a 0.5 Log2 upregulation in control LEC vs knock-out LEC or genes that had at least a 0.5 Log2 upregulation in knock-out LECs vs. control LECs were utilized for over-representation analysis. False discovery rate (FDR)-adjusted *P* values of less than 0.05 were considered statistically significant.

### Statistical analysis and data reporting

Data were assessed and analyzed using appropriate statistical methods. The normality of data where appropriate was assessed using the Kolmogorov–Smirnov test. GraphPad Prism v.9 was used for all statistical analysis, unless otherwise indicated. No statistical methods were used to determine sample size. Unless otherwise stated, the experiments were not randomized. The investigators were blinded during all 5FU and DT experiments quantifications.

---

**Supplementary Figure 1.**
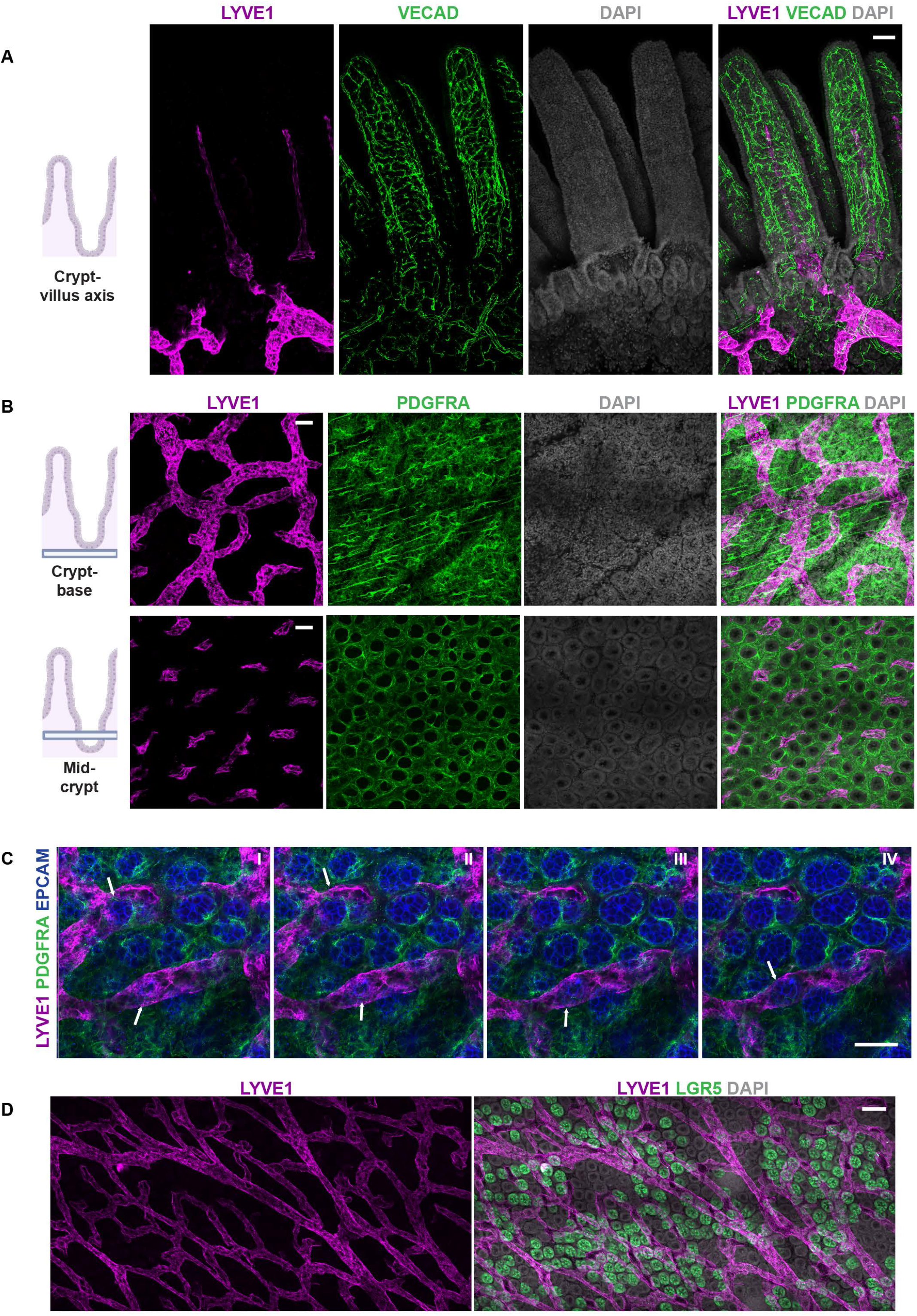
LECs are in proximity to crypt epithelial cells. **A)** Whole mount tissue imaging, from sagittal view, of LYVE1 and VECAD staining showing the localization of lymphatic and blood vessels within the crypt-villus axis in the small intestine. Scale bar = 50 μm. **B)** Whole mount tissue imaging, from transverse view, of LYVE1 and PDGFRA staining showing the proximity of LECs to the base and the mid-part of crypts. Scale bar = 50 μm. **C)** Consecutive orthogonal slices of LECs (LYVE1^+^) and epithelial cells (EPCAM^+^) from the base of the crypts to mid-crypt area. White arrows represent point of contact between LECs and epithelial crypts cells. Scale bar = 50 μm. **D)** Whole mount imaging in *Lgr5*^DTR-EGFP^ mice, from transverse view, of LYVE1 and *Lgr5*-GFP crypt cells. Scale bar = 100 μm.

**Supplementary Figure 2.**
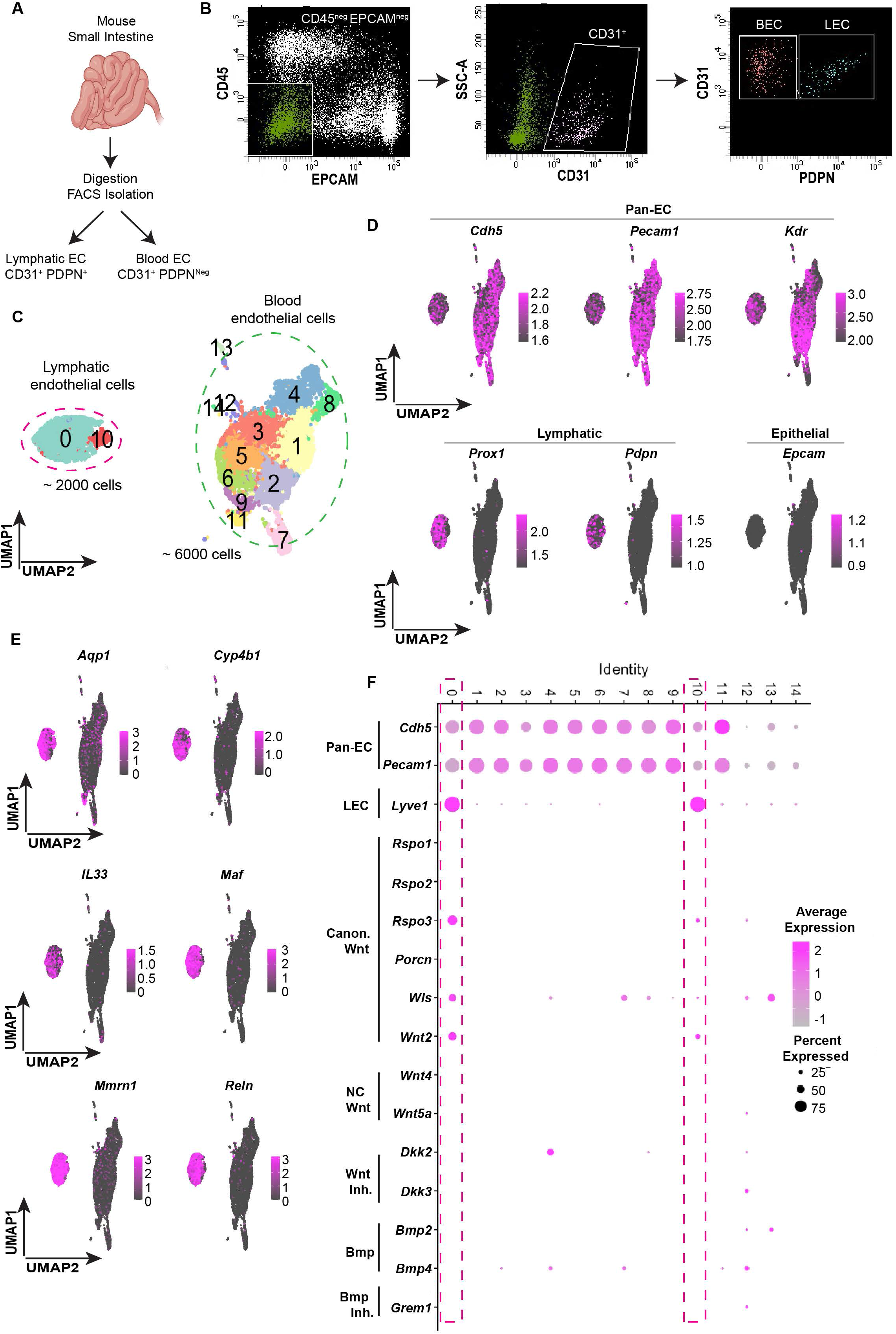
LEC scRNAseq data reveal expression of Wnt signaling molecules. **A)** Small intestines from 4 wild-type mice were digested, and LECs and BECs were isolated by sorting CD31^+^, PDPN^+^ and CD31^+^, PDPN^Neg^ cells, respectively. **B)** Gating strategy to isolate LECs and BECs. First, we selected CD45^Neg^ and EPCAM^Neg^ cells to exclude immune and epithelial cells, then CD31^+^ endothelial cells were isolated, and finally LECs and BECs were distinguished as PDPN^+^ and PDPN^Neg^, respectively. **C)** UMAP plot of all ECs showing that LECs group in 2 clusters (cluster 0 and 10) and BECs group in 12 different clusters. **D)** scRNAseq expression of markers of pan-endothelial, lymphatic, and epithelial cells validating the identity of LECs and BECs. **E)** scRNAseq expression of unique LEC markers compared to BECs. **F)**Dotplot of relative gene expression in the 14 EC clusters of canonical Wnt ligands and modulators, non-canonical Wnt ligands, Wnt inhibitors, Bmp ligands and inhibitors.

**Supplementary Figure 3.**
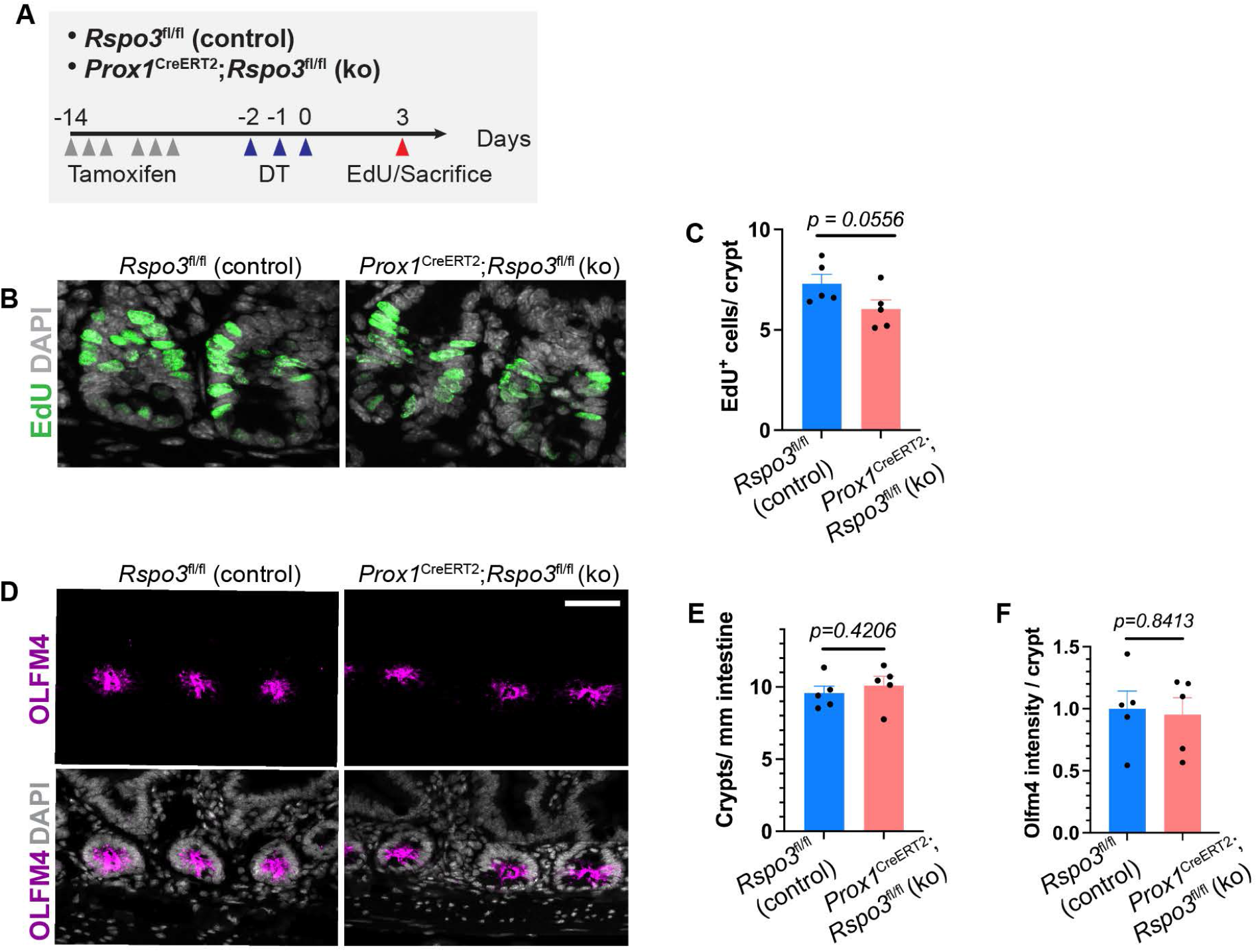
LEC derived *Rspo3* is dispensable for intestinal recovery after ablation of *Lgr* 5+ cells. **A)** Tamoxifen induced *Rspo3* knock-out (ko) and control (ctrl) mice were treated with DT for 3 consecutive days to remove *Lgr5*^+^ ISC. Mice were injected with EdU before sacrifice 3 days after the last DT injection. **B)** Representative images of EdU staining and quantification of EdU^+^ cells per crypt and **(C)** of ctrl and ko mice after DT treatment (n=5 mice per group). **D)** Representative images of OLFM4 staining (scale bar = 50 μm), **(E)** quantification of OLFM4^+^ crypt number (scale bar = 50 μm), **(F)** OLFM4 staining intensity per crypt in control and *Rspo3* knock-out mice after DT treatment. Data are mean +/− s.e.m., Mann Whitney U-test, two-sided.

**Supplementary Figure 4.**
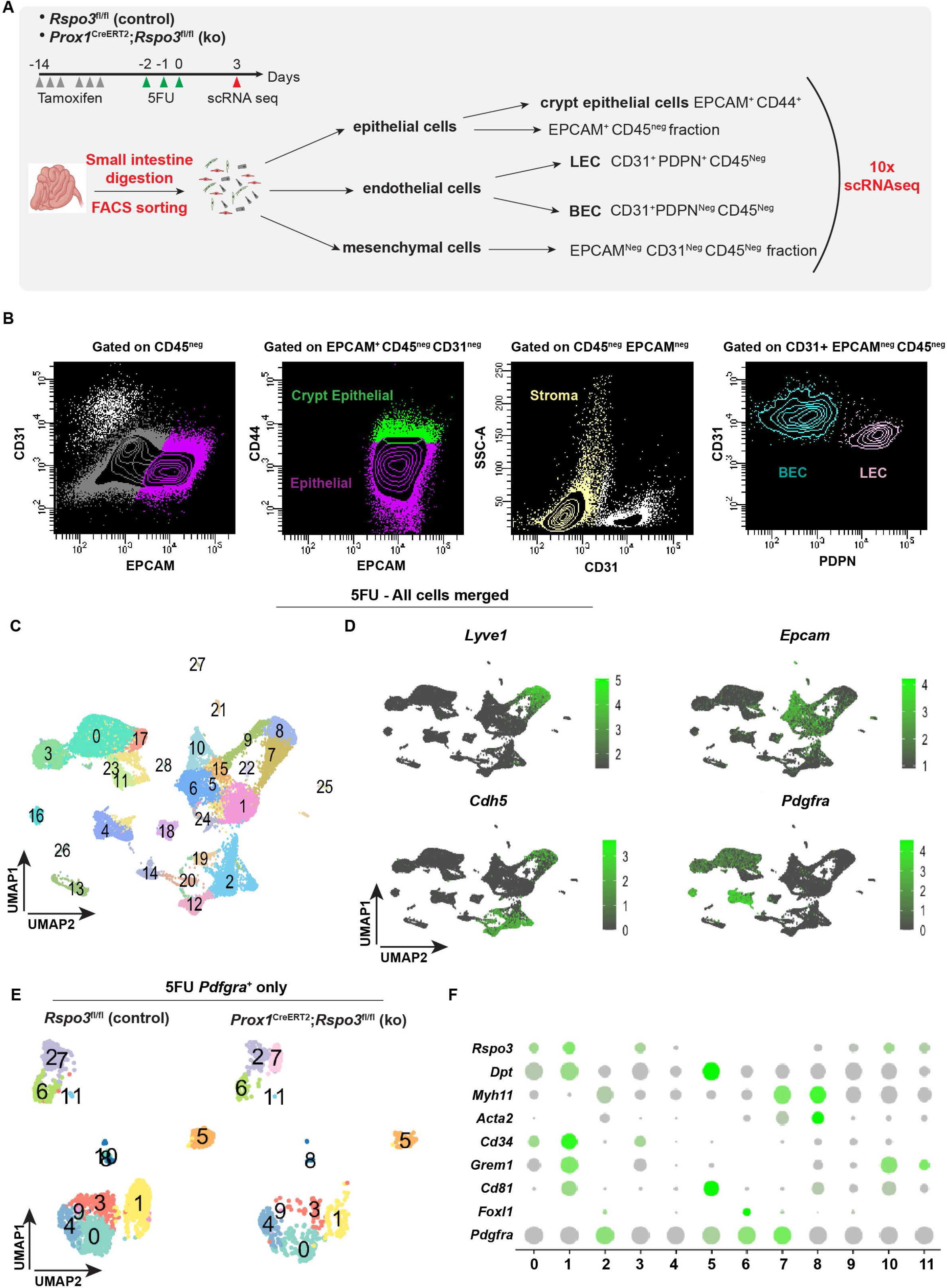
scRNAseq data after 5FU injury for LEC, BEC, crypt epithelial cells and mesenchymal cells. **A)** Schema of scRNAseq experiment. Tamoxifen treated control and knock-out mice were administered 5FU on 3 consecutive days. Crypt epithelial cells (EPCAM^+^, CD44^+^), LECs (CD31^+^, PDPN^+^), BECs (CD31^+^, PDPN^Neg^) and other stromal cells (EPCAM^Neg^, CD31^Neg^) were enriched by FACS, and the samples submitted for scRNAseq analysis. Data are from 2 independent biological experiments. Each biological experiment included 2 control and 2 knock-out mice. **B)** Gating strategy to isolate crypt epithelial cells, LECs, BECs, epithelial cells and other stromal cells. For crypt epithelial cells, we first selected double CD45^Neg^ and CD31^Neg^ cells to exclude immune and endothelial cells, respectively. Next, EPCAM^+^, CD44^+^ crypt epithelial cells were isolated. For ECs, we selected cells that were double negative EPCAM^Neg^, CD45^Neg^ (to exclude epithelial and immune cells, respectively) and positive for CD31. LECs and BECs were distinguished as PDPN^+^ and PDPN^Neg^, respectively. For other stromal cells, we selected triple negative cells as EPCAM^Neg^, CD45^Neg^, CD31^Neg^. **C)** UMAP plot of combined cells from all samples, which group into 28 distinct clusters. **D)** scRNAseq expression of pan endothelial (*Vecad*), lymphatic (*Lyve1*), epithelial (*Epcam*), and mesenchymal (*Pdgfra*) cell markers. **E)** Mesenchymal cells were isolated as *Pdgfra*^*+*^ cells and re-clustered. UMAP of *Pdgfra*^*+*^ cells showing they group in 12 unique clusters. **F)** Dotplot of markers present in *Pdgfra*^*+*^ cells identifying telocytes as *Foxl1*^*+*^, trophocytes as *Cd81*^*+*^, *Grem1*^*+*^ cells, pericryptal stroma cells as *Cd34*^*+*^, fibroblasts as *Dpt*^*+*^ and myofibroblasts as *Myh11*^*+*^.

## References

Abel, E., Ekman, T., Warnhammar, E., Hultborn, R., Jennische, E., and Lange, S. (2005). Early disturbance of microvascular function precedes chemotherapy-induced intestinal injury. Dig Dis Sci 50, 1729–1733. 10.1007/s10620-005-2926-9.

Abu El-Asrar, A.M., Ahmad, A., Siddiquei, M.M., De Zutter, A., Allegaert, E., Gikandi, P.W., De Hertogh, G., Van Damme, J., Opdenakker, G., and Struyf, S. (2019). The Proinflammatory and Proangiogenic Macrophage Migration Inhibitory Factor Is a Potential Regulator in Proliferative Diabetic Retinopathy. Front Immunol 10, 2752. 10.3389/fimmu.2019.02752.

Aoki, R., Shoshkes-Carmel, M., Gao, N., Shin, S., May, C.L., Golson, M.L., Zahm, A.M., Ray, M., Wiser, C.L., Wright, C.V., and Kaestner, K.H. (2016). Foxl1-expressing mesenchymal cells constitute the intestinal stem cell niche. Cell Mol Gastroenterol Hepatol 2, 175–188. 10.1016/j.jcmgh.2015.12.004.

Augustin, H.G., and Koh, G.Y. (2017). Organotypic vasculature: From descriptive heterogeneity to functional pathophysiology. Science 357. 10.1126/science.aal2379.

Barker, N., Es, J.H.v., Kuipers, J., Kujala, P., Born, M.v.d., Cozijnsen, M., Haegebarth, A., Korving, J., Begthel, H., Peters, P.J., and Clevers, H. (2007). Identification of stem cells in small intestine and colon by marker gene Lgr5. Nature 449, 1003–1007. doi:10.1038/nature06196.

Bayoumi, A.S., Teoh, J.P., Aonuma, T., Yuan, Z., Ruan, X., Tang, Y., Su, H., Weintraub, N.L., and Kim, I.M. (2017). MicroRNA-532 protects the heart in acute myocardial infarction, and represses prss23, a positive regulator of endothelial-to-mesenchymal transition. Cardiovasc Res 113, 1603–1614. 10.1093/cvr/cvx132.

Bernier-Latmani, J., Cisarovsky, C., Demir, C.S., Bruand, M., Jaquet, M., Davanture, S., Ragusa, S., Siegert, S., Dormond, O., Benedito, R., et al. (2015). DLL4 promotes continuous adult intestinal lacteal regeneration and dietary fat transport. 10.1172/JCI82045.

Bernier-Latmani, J., and Petrova, T.V. (2017). Intestinal lymphatic vasculature: structure, mechanisms and functions. Nat Rev Gastroenterol Hepatol 14, 510–526. 10.1038/nrgastro.2017.79.

Beumer, J., and Clevers, H. (2020). Cell fate specification and differentiation in the adult mammalian intestine. Nature Reviews Molecular Cell Biology 22, 39–53. doi:10.1038/s41580-020-0278-0.

Butler, A., Hoffman, P., Smibert, P., Papalexi, E., and Satija, R. (2018). Integrating single-cell transcriptomic data across different conditions, technologies, and species. Nat Biotechnol 36, 411–420. 10.1038/nbt.4096.

Butler, J.M., Nolan, D.J., Vertes, E.L., Varnum-Finney, B., Kobayashi, H., Hooper, A.T., Seandel, M., Shido, K., White, I.A., Kobayashi, M., et al. (2010). Endothelial cells are essential for the self-renewal and repopulation of Notch-dependent hematopoietic stem cells. Cell Stem Cell 6, 251–264. 10.1016/j.stem.2010.02.001.

Cifarelli, V., and Eichmann, A. (2019). The Intestinal Lymphatic System: Functions and Metabolic Implications. Cell Mol Gastroenterol Hepatol 7, 503–513. 10.1016/j.jcmgh.2018.12.002.

de Lau, W., Barker, N., Low, T.Y., Koo, B.-K., Li, V.S.W., Teunissen, H., Kujala, P., Haegebarth, A., Peters, P.J., van de Wetering, M., et al. (2011). Lgr5 homologues associate with Wnt receptors and mediate R-spondin signalling. Nature 476, 293–297. doi:10.1038/nature10337.

Degirmenci, B., Valenta, T., Dimitrieva, S., Hausmann, G., and Basler, K. (2018). GLI1-expressing mesenchymal cells form the essential Wnt-secreting niche for colon stem cells. Nature 558, 449–453. 10.1038/s41586-018-0190-3.

Ding, B.S., Nolan, D.J., Butler, J.M., James, D., Babazadeh, A.O., Rosenwaks, Z., Mittal, V., Kobayashi, H., Shido, K., Lyden, D., et al. (2010). Inductive angiocrine signals from sinusoidal endothelium are required for liver regeneration. Nature 468, 310–315. 10.1038/nature09493.

Ding, B.S., Nolan, D.J., Guo, P., Babazadeh, A.O., Cao, Z., Rosenwaks, Z., Crystal, R.G., Simons, M., Sato, T.N., Worgall, S., et al. (2011). Endothelial-derived angiocrine signals induce and sustain regenerative lung alveolarization. Cell 147, 539–553. 10.1016/j.cell.2011.10.003.

Ding, L., Saunders, T.L., Enikolopov, G., and Morrison, S.J. (2012). Endothelial and perivascular cells maintain haematopoietic stem cells. Nature 481, 457–462. 10.1038/nature10783.

Forman, R.A., deSchoolmeester, M.L., Hurst, R.J., Wright, S.H., Pemberton, A.D., and Else, K.J. (2012). The goblet cell is the cellular source of the anti-microbial angiogenin 4 in the large intestine post Trichuris muris infection. PLoS One 7, e42248. 10.1371/journal.pone.0042248.

Gehart, H., and Clevers, H. (2019). Tales from the crypt: new insights into intestinal stem cells. Nat Rev Gastroenterol Hepatol 16, 19–34. 10.1038/s41575-018-0081-y.

Glinka, A., Dolde, C., Kirsch, N., Huang, Y.L., Kazanskaya, O., Ingelfinger, D., Boutros, M., Cruciat, C.M., and Niehrs, C. (2011). LGR4 and LGR5 are R-spondin receptors mediating Wnt/β-catenin and Wnt/PCP signalling. EMBO Rep 12, 1055–1061. 10.1038/embor.2011.175.

Goto, N., Imada, S., Deshpande, V., and Yilmaz, Ö.H. (2022). Lymphatics constitute a novel component of the intestinal stem cell niche. BioRxiv 10.1101/2022.01.28.478205.

Greicius, G., Kabiri, Z., Sigmundsson, K., Liang, C., Bunte, R., Singh, M.K., and Virshup, D.M. (2018). pericryptal stromal cells are the critical source of Wnts and RSPO3 for murine intestinal stem cells in vivo. Proc Natl Acad Sci U S A 115, E3173–E3181. 10.1073/pnas.1713510115.

Gur-Cohen, S., Yang, H., Baksh, S.C., Miao, Y., Levorse, J., Kataru, R.P., Liu, X., Cruz-Racelis, J.d.l., Mehrara, B.J., and Fuchs, E. (2019). Stem cell–driven lymphatic remodeling coordinates tissue regeneration. 10.1126/science.aay4509.

Haber, A.L., Biton, M., Rogel, N., Herbst, R.H., Shekhar, K., Smillie, C., Burgin, G., Delorey, T.M., Howitt, M.R., Katz, Y., et al. (2017). A single-cell survey of the small intestinal epithelium. Nature 551, 333–339. doi:10.1038/nature24489.

Hao, H.-X., Xie, Y., Zhang, Y., Charlat, O., Oster, E., Avello, M., Lei, H., Mickanin, C., Liu, D., Ruffner, H., et al. (2012). ZNRF3 promotes Wnt receptor turnover in an R-spondin-sensitive manner. Nature 485, 195–200. doi:10.1038/nature11019.

Harnack, C., Berger, H., Antanaviciute, A., Vidal, R., Sauer, S., Simmons, A., Meyer, T.F., and Sigal, M. (2019). R-spondin 3 promotes stem cell recovery and epithelial regeneration in the colon. Nat Commun 10, 4368. 10.1038/s41467-019-12349-5.

Hooper, L.V., Stappenbeck, T.S., Hong, C.V., and Gordon, J.I. (2003). Angiogenins: a new class of microbicidal proteins involved in innate immunity. Nat Immunol 4, 269–273. 10.1038/ni888.

Hu, J., Srivastava, K., Wieland, M., Runge, A., Mogler, C., Besemfelder, E., Terhardt, D., Vogel, M.J., Cao, L., Korn, C., et al. (2014). Endothelial cell-derived angiopoietin-2 controls liver regeneration as a spatiotemporal rheostat. Science 343, 416–419. 10.1126/science.1244880.

Kabiri, Z. (2014). Stroma provides an intestinal stem cell niche in the absence of epithelial Wnts. http://dx.doi.org/10.1242/dev.104976.

Kalucka, J., de Rooij, L.P.M.H., Goveia, J., Rohlenova, K., Dumas, S.J., Meta, E., Conchinha, N.V., Taverna, F., Teuwen, L.A., Veys, K., et al. (2020). Single-Cell Transcriptome Atlas of Murine Endothelial Cells. Cell 180, 764–779.e720. 10.1016/j.cell.2020.01.015.

Kinchen, J., Chen, H.H., Parikh, K., Antanaviciute, A., Jagielowicz, M., Fawkner-Corbett, D., Ashley, N., Cubitt, L., Mellado-Gomez, E., Attar, M., et al. (2018). Structural Remodeling of the Human Colonic Mesenchyme in Inflammatory Bowel Disease. Cell 175, 372–386.e317. 10.1016/j.cell.2018.08.067.

Kondo, A., Department of Genetics, P.S.o.M., University of Pennsylvania, Philadelphia, PA 19104,◻USA, Kaestner, K.H., and Department of Genetics, P.S.o.M., University of Pennsylvania, Philadelphia, PA 19104,◻USA (2019). Emerging diverse roles of telocytes. Development 146. 10.1242/dev.175018.

Li, X., Liu, B., Xiao, J., Yuan, Y., Ma, J., and Zhang, Y. (2011). Roles of VEGF-C and Smad4 in the lymphangiogenesis, lymphatic metastasis, and prognosis in colon cancer. J Gastrointest Surg 15, 2001–2010. 10.1007/s11605-011-1627-2.

Liao, Y., Wang, J., Jaehnig, E.J., Shi, Z., and Zhang, B. (2019). WebGestalt 2019: gene set analysis toolkit with revamped UIs and APIs. Nucleic Acids Res 47, W199–W205. 10.1093/nar/gkz401.

Liu, X., De la Cruz, E., Gu, X., Balint, L., Oxendine-Burns, M., Terrones, T., Ma, W., Kuo, H.-H., Lantz, C., Bansal, T., et al. (2020). Lymphoangiocrine signals promote cardiac growth and repair. Nature 588, 705–711. doi:10.1038/s41586-020-2998-x.

Maleszewska, M., Moonen, J.R., Huijkman, N., van de Sluis, B., Krenning, G., and Harmsen, M.C. (2013). IL-1β and TGFβ2 synergistically induce endothelial to mesenchymal transition in an NFκB-dependent manner. Immunobiology 218, 443–454. 10.1016/j.imbio.2012.05.026.

Matsumoto, K., Mitani, T.T., Horiguchi, S.A., Kaneshiro, J., Murakami, T.C., Mano, T., Fujishima, H., Konno, A., Watanabe, T.M., Hirai, H., and Ueda, H.R. (2019). Advanced CUBIC tissue clearing for whole-organ cell profiling. Nature Protocols 14, 3506–3537. doi:10.1038/s41596-019-0240-9.

McCarthy, N., Kraiczy, J., and Shivdasani, R.A. (2020a). Cellular and molecular architecture of the intestinal stem cell niche. Nature Cell Biology 22, 1033–1041. doi:10.1038/s41556-020-0567-z.

McCarthy, N., Manieri, E., Storm, E.E., Saadatpour, A., Luoma, A.M., Kapoor, V.N., Madha, S., Gaynor, L.T., Cox, C., Keerthivasan, S., et al. (2020b). Distinct Mesenchymal Cell Populations Generate the Essential Intestinal BMP Signaling Gradient. Cell Stem Cell 26, 391–402.e395. 10.1016/j.stem.2020.01.008.

Nusse, Y.M., Savage, A.K., Marangoni, P., Rosendahl-Huber, A.K.M., Landman, T.A., de Sauvage, F.J., Locksley, R.M., and Klein, O.D. (2018). Parasitic helminths induce fetal-like reversion in the intestinal stem cell niche. Nature 559, 109–113. 10.1038/s41586-018-0257-1.

Ogasawara, R., Hashimoto, D., Kimura, S., Hayase, E., Ara, T., Takahashi, S., Ohigashi, H., Yoshioka, K., Tateno, T., Yokoyama, E., et al. (2018). Intestinal Lymphatic Endothelial Cells Produce R-Spondin3. Scientific Reports 8, 1–9. doi:10.1038/s41598-018-29100-7.

Palikuqi, B., Rispal, J., and Klein, O. (2021). Good Neighbors: The Niche that Fine Tunes Mammalian Intestinal Regeneration. 10.1101/cshperspect.a040865.

Paris, F., Fuks, Z., Kang, A., Capodieci, P., Juan, G., Ehleiter, D., Haimovitz-Friedman, A., Cordon-Cardo, C., and Kolesnick, R. (2001). Endothelial apoptosis as the primary lesion initiating intestinal radiation damage in mice. Science 293, 293–297. 10.1126/science.1060191.

Rafii, S., Butler, J.M., and Ding, B.S. (2016). Angiocrine functions of organ-specific endothelial cells. Nature 529, 316–325. 10.1038/nature17040.

Rehal, S., Stephens, M., Roizes, S., Liao, S., and von der Weid, P.Y. (2018). Acute small intestinal inflammation results in persistent lymphatic alterations. Am J Physiol Gastrointest Liver Physiol 314, G408–G417. 10.1152/ajpgi.00340.2017.

Ruffell, D., Mourkioti, F., Gambardella, A., Kirstetter, P., Lopez, R.G., Rosenthal, N., and Nerlov, C. (2009). A CREB-C/EBPbeta cascade induces M2 macrophage-specific gene expression and promotes muscle injury repair. Proc Natl Acad Sci U S A 106, 17475–17480. 10.1073/pnas.0908641106.

Sato, T., Vries, R.G., Snippert, H.J., van de Wetering, M., Barker, N., Stange, D.E., van Es, J.H., Abo, A., Kujala, P., Peters, P.J., and Clevers, H. (2009). Single Lgr5 stem cells build crypt-villus structures in vitro without a mesenchymal niche. Nature 459, 262–265. 10.1038/nature07935.

Scholz, B., Korn, C., Wojtarowicz, J., Mogler, C., Augustin, I., Boutros, M., Niehrs, C., and Augustin, H.G. (2016). Endothelial RSPO3 Controls Vascular Stability and Pruning through Non-canonical WNT/Ca(2+)/NFAT Signaling. Dev Cell 36, 79–93. 10.1016/j.devcel.2015.12.015.

Shoshkes-Carmel, M., Wang, Y.J., Wangensteen, K.J., Tóth, B., Kondo, A., Massasa, E.E., Itzkovitz, S., and Kaestner, K.H. (2018). Subepithelial telocytes are an important source of Wnts that supports intestinal crypts. Nature 557, 242–246. doi:10.1038/s41586-018-0084-4.

Storm, E.E., Durinck, S., de Sousa e Melo, F., Tremayne, J., Kljavin, N., Tan, C., Ye, X., Chiu, C., Pham, T., Hongo, J.-A., et al. (2015). Targeting PTPRK-RSPO3 colon tumours promotes differentiation and loss of stem-cell function. Nature 529, 97–100. doi:10.1038/nature16466.

Stzepourginski, I., Nigro, G., Jacob, J.M., Dulauroy, S., Sansonetti, P.J., Eberl, G., and Peduto, L. (2017). CD34+ mesenchymal cells are a major component of the intestinal stem cells niche at homeostasis and after injury. Proc Natl Acad Sci U S A 114, E506–E513. 10.1073/pnas.1620059114.

Tacconi, C., Correale, C., Gandelli, A., Spinelli, A., Dejana, E., D’Alessio, S., and Danese, S. (2015). Vascular endothelial growth factor C disrupts the endothelial lymphatic barrier to promote colorectal cancer invasion. Gastroenterology 148, 1438–1451.e1438. 10.1053/j.gastro.2015.03.005.

Tian, H., Biehs, B., Warming, S., Leong, K.G., Rangell, L., Klein, O.D., and de Sauvage, F.J. (2011). A reserve stem cell population in small intestine renders Lgr5-positive cells dispensable. Nature 478, 255–259. 10.1038/nature10408.

Valenta, T., Degirmenci, B., Moor, A.E., Herr, P., Zimmerli, D., Moor, M.B., Hausmann, G., Cantù, C., Aguet, M., and Basler, K. (2016). Wnt Ligands Secreted by Subepithelial Mesenchymal Cells Are Essential for the Survival of Intestinal Stem Cells and Gut Homeostasis. Cell Rep 15, 911–918. 10.1016/j.celrep.2016.03.088.

Wertheimer, T., Velardi, E., Tsai, J., Cooper, K., Xiao, S., Kloss, C.C., Ottmuller, K.J., Mokhtari, Z., Brede, C., deRoos, P., et al. (2018). Production of BMP4 by endothelial cells is crucial for endogenous thymic regeneration. Sci Immunol 3. 10.1126/sciimmunol.aal2736.

Yan, K.S., Janda, C.Y., Chang, J., Zheng, G.X.Y., Larkin, K.A., Luca, V.C., Chia, L.A., Mah, A.T., Han, A., Terry, J.M., et al. (2017). Non-equivalence of Wnt and R-spondin ligands during Lgr5. Nature 545, 238–242. 10.1038/nature22313.

Yang, X., Kim, J.-d., Gu, Q., Yan, Q., Astin, J., Crosier, P.S., Yu, P., Rockson, S.G., and Fang, L. (2020). AIBP-CAV1-VEGFR3 axis dictates lymphatic cell fate and controls lymphangiogenesis. 10.1101/2020.10.16.342998.

Yoshimatsu, Y., Kimuro, S., Pauty, J., Takagaki, K., Nomiyama, S., Inagawa, A., Maeda, K., Podyma-Inoue, K.A., Kajiya, K., Matsunaga, Y.T., and Watabe, T. (2020). TGF-beta and TNF-alpha cooperatively induce mesenchymal transition of lymphatic endothelial cells via activation of Activin signals. PLoS One 15, e0232356. 10.1371/journal.pone.0232356.

Zheng, W., Rosenstiel, P., Huse, K., Sina, C., Valentonyte, R., Mah, N., Zeitlmann, L., Grosse, J., Ruf, N., Nürnberg, P., et al. (2006). Evaluation of AGR2 and AGR3 as candidate genes for inflammatory bowel disease. Genes Immun 7, 11–18. 10.1038/sj.gene.6364263.

